# Mid-summer snow-free albedo across the Arctic tundra was mostly stable or increased over the past two decades

**DOI:** 10.1101/2022.07.11.499587

**Authors:** Elena Plekhanova, Jin-Soo Kim, Jacqueline Oehri, Angela Erb, Crystal Schaaf, Gabriela Schaepman-Strub

## Abstract

Arctic vegetation changes, such as increasing shrub-cover, are expected to accelerate climate warming through increased absorption of incoming radiation and corresponding decrease in summer shortwave albedo. Here we analyze mid-summer shortwave land-surface albedo and its change across the pan-Arctic region based on MODIS satellite observations over the past two decades (2000-2021). In contrast to expectations, we show that terrestrial mid-summer shortwave albedo has not significantly changed in 82% of the pan-Arctic region, while 14% show an increase and 4% a decrease. By analyzing the visible and near-/shortwave-infrared range separately, we demonstrate that the slight increase arises from an albedo increase in the near-/shortwave-infrared range domain while being partly compensated by a decrease in visible albedo. A similar response was found across different tundra vegetation types. We argue that this increase in reflectance is typical with increasing biomass as a result of increased multiple reflection in the canopy. However, CMIP6 global climate model albedo predictions showed the opposite sign and different spatial patterns of snow-free summer albedo change compared to satellite-derived results. We suggest that a more sophisticated vegetation parametrization can reduce this discrepancy, and provide albedo estimates per vegetation type.

## 1. Introduction

Recent climate change has induced multiple biogeophysical changes in Arctic ecosystems, including shifts in snow phenology and vegetation composition. Such land surface changes can further amplify or reduce climate warming by altering the surface reflectance - the albedo. Albedo is a key variable in the land surface components of climate and Earth system models. Therefore, to properly account for the land surface feedback under amplified Arctic warming, we need to quantify albedo change, understand its mechanisms, and adequately include it in climate models.

The largest and most studied terrestrial Arctic albedo change occurs in the shoulder seasons when highly reflective snow gives way to much darker land covers (vegetation, water, soil), or vice versa. Indeed, recent satellite observations show an advance in snow-melt date and a delay in freezing, which has increased the snow-free period by about 4 days per decade (Chen *et al*., 2015), and has created a strong positive albedo-climate feedback. However, with the lengthening of the snow-free periods in the high latitudes, the much less studied snow-free summer albedo changes due to vegetation development become increasingly important (Chapin *et al*., 2005).

In the snow-free period, terrestrial Arctic land surface albedo is mostly driven by water cover and vegetation cover and structure (Oehri, *et al*., in review; Juszak *et al*., 2017; Webb *et al*., 2021). Surface water usually absorbs more shortwave radiation compared to other land cover types, so an increase in water cover would decrease the albedo. However, in recent decades the signal is mixed: the discharge of main rivers has increased in the Eurasian river basins and has decreased in the North American basins (Box *et al*., 2019; Lin *et al*., 2022), while lake area has mostly decreased (Smith *et al*., 2005; Finger Higgens *et al*., 2019). Besides water, the Arctic tundra is very heterogeneous and covered by a variety of vegetation types. The distribution of 5 main vegetation types (prostrate and erect shrubs, graminoids, wetlands, and barrens (Walker *et al*., 2005, Supplementary Table 1)) is captured by the Circumpolar Arctic Vegetation Map (CAVM, Raynolds *et al*., 2019). This framework has been found useful for many applications including the prediction of surface energy fluxes (Oehri, *et al*., in review).

The most widely reported vegetation change in the Arctic is a greening, detected from satellite observations as an increase in the Normalized Difference Vegetation Index (NDVI), associated with biomass increase (Bhatt *et al*., 2017; Beamish *et al*., 2020). More recent studies revealed complex and scale-dependent greening and browning trends (Myers-Smith *et al*., 2020, Ju & Masek, 2016). The increase in biomass is explained by shrubs becoming taller (Bjorkman *et al*., 2020) and more widespread (Sturm, Racine and Tape, 2001; Myers-Smith *et al*., 2011; Martin *et al*., 2017), while lichens and mosses have decreased in abundance (CAFF, 2021). Earlier studies have suggested that this shrubification, together with a treeline advance, would decrease summer albedo (Thompson *et al*., 2004; Chapin *et al*., 2005) and hence cause a positive feedback with Arctic warming. Contrary to these expectations, recent observation-based studies have reported an increase in summer albedo. High-latitude July albedo in certain areas has increased by 3-9% per year (Webb *et al*., 2021; Yu and Leng, 2022), which resulted in a negative surface-albedo feedback of −0.91 W/(m2 · K) (Alessandri *et al*., 2021), and a 5% albedo increase per degree of warming (Yu, Leng and Python, 2022). However, it remains unclear how much of the pan-Arctic area has actually experienced significant albedo trends and what the reasons are for the reported unexpected observations.

Most of the studies investigate the broadband shortwave albedo as a spectral integral quantity. However, the total incoming solar radiation (broadband shortwave domain, 300-5000 nm) includes a) a visible domain (VIS, 300-700nm) and b) a near-infrared to shortwave infrared domain (NIR+SWIR, 700-5000 nm). Reflectance of vegetation and other land cover types is very different between those two spectral domains. Hence it is important to separate VIS and NIR+SWIR for energy budget applications, such as climate modeling (Roesch *et al*., 2002).

In climate models, the changes in land cover and its albedo are represented in the land surface model (LSM). Several studies compared modeled albedo with satellite observations to ensure concordance (Crook and Forster, 2014; Levine and Boos, 2017; Thackeray, Fletcher and Derksen, 2019). Most land surface albedo studies however have focused on the more extreme shoulder season variability (Crook and Forster, 2014; Loranty *et al*., 2014), oftenly entirely excluding high latitudes (Jian et al., 2020), and have not compared albedo trends across decades for the pan-Arctic tundra region.

Hence, the goal of this study is 1) to assess and locate trends in pan-Arctic snow-free mid-summer albedo across spectral domains and tundra vegetation types during the past 22 years and 2) to quantify the differences between the mid-summer albedo values and trends observed by satellites and predicted by climate models. We use snow-free mid-summer (15th of July) MODerate resolution Imaging Spectroradiometer (MODIS) albedo products (Schaaf *et al*., 2002; Wang *et al*., 2012, 2018) from 2000 to 2022 to quantify albedo change per year across the CAVM extent and compare it with albedo change derived from LSMs from the latest climate model intercomparison project (i.e., CMIP6) (Eyring *et al*., 2016). Finally, we discuss how our findings contribute to a better understanding of mechanisms of Arctic albedo change and its application in future predictions of change in energy budget under Arctic amplification.

## 2. Results

Here, we first show pan-Arctic snow-free land surface albedo trends across the broadband shortwave, VIS, and NIR+SWIR spectral ranges. We then compare albedo change across vegetation types. Finally, we highlight how albedo and its changes are incorporated in current land surface models (LSM) and compare these LSM estimates with results from MODIS-based observations.

### 2.1. Snow-free albedo trends across the Arctic

Based on 22 years (2000-2021) of MODIS data, we estimated mid-summer snow-free albedo trends per year (via Mann-Kendall regression, Theil–Sen slope estimator (Sen, 1968), Kendall’s tau significance statistic, see Methods, Fig. 1 a,d). We found that 82% of the pan-Arctic land area did not show any significant trend in shortwave albedo. The remaining 18% (∼ 8.4 ×10^5^ km^2^) of the Arctic landscape did change significantly with 14% showing a positive trend and 4% a negative trend. Most of these trends were below 0.001 per year (1% change per year), with a total median significant change of 0.014 over the past 22 years.

**Fig. 1.**
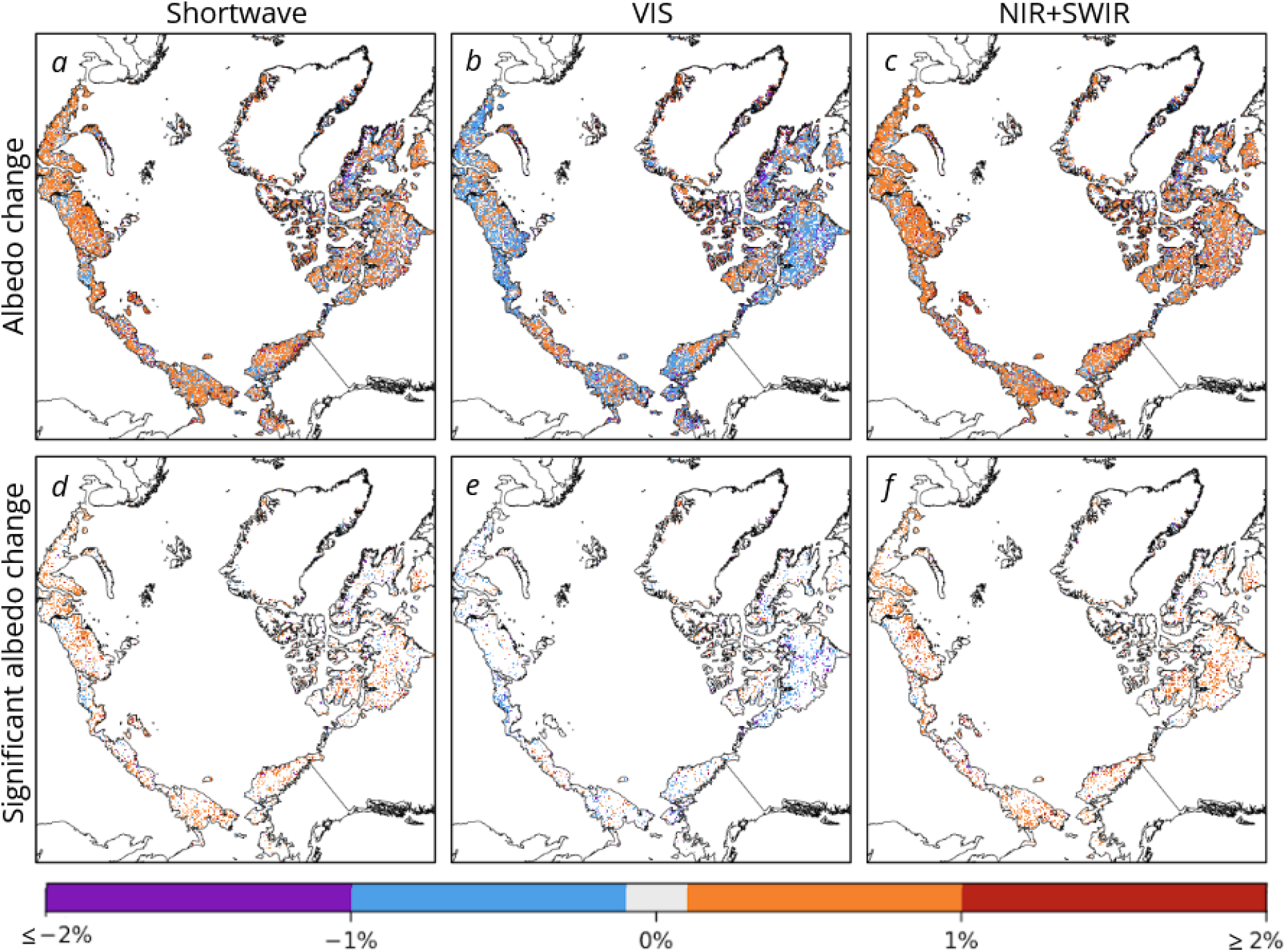
Pan-Arctic mid-summer land surface albedo trends (2000-2021) in the broadband shortwave (300-5000nm) (a,d), visible (300-700nm) (b,e) and near- to shortwave infrared (NIR+SWIR) (700-5000 nm) (c,f) wavelength range of the spectrum. The upper row (a-c) shows estimated percentage of albedo change per year (based on Theil–Sen’s slope), and the bottom row (d-f) shows only areas with significant (p-value < 0.05) changes.

### 2.2 Why does the shortwave albedo increase?

To answer this question, we investigated the change in the VIS and NIR+SWIR part of the shortwave range separately (see Fig.1 b,c,e,f). In the VIS range, 13% of slopes were significant: 3% positive and 10% negative. In the NIR+SWIR range, 19% of slopes were significant: 16% positive and 3% negative. Overlap between significant VIS and NIR+SWIR trends occurred in 4% of the area, between VIS and shortwave in 4%, while 12% of the area showed concurrent significant trends between the NIR+SWIR and shortwave albedo (Supplementary Fig. 1). The total median significant change integrated over the past 22 years was −13.2% (−0.007) in the visible range, and 11.3% (0.025) in the NIR+SWIR. Hence, the overall shortwave albedo trend is slightly increasing because the positive trend in NIR+SWIR is stronger than the negative trend in VIS, while 12% of area (67% of significant shortwave trends) shows significant change in both shortwave and NIR+SWIR domains.

### 2.3. Change in albedo of different vegetation types

We also analyzed the significant shortwave, VIS and NIR+SWIR albedo slopes over structurally different Arctic vegetation types (see Fig. 2). There is a substantial difference between the albedo of different vegetation types, especially when separated into VIS and NIR+SWIR domains (Fig. 2). Consistent with the previous result, albedo in the broadband shortwave and the NIR+SWIR domains, on average, increased, but slightly decreased in the VIS range.

**Fig. 2.**
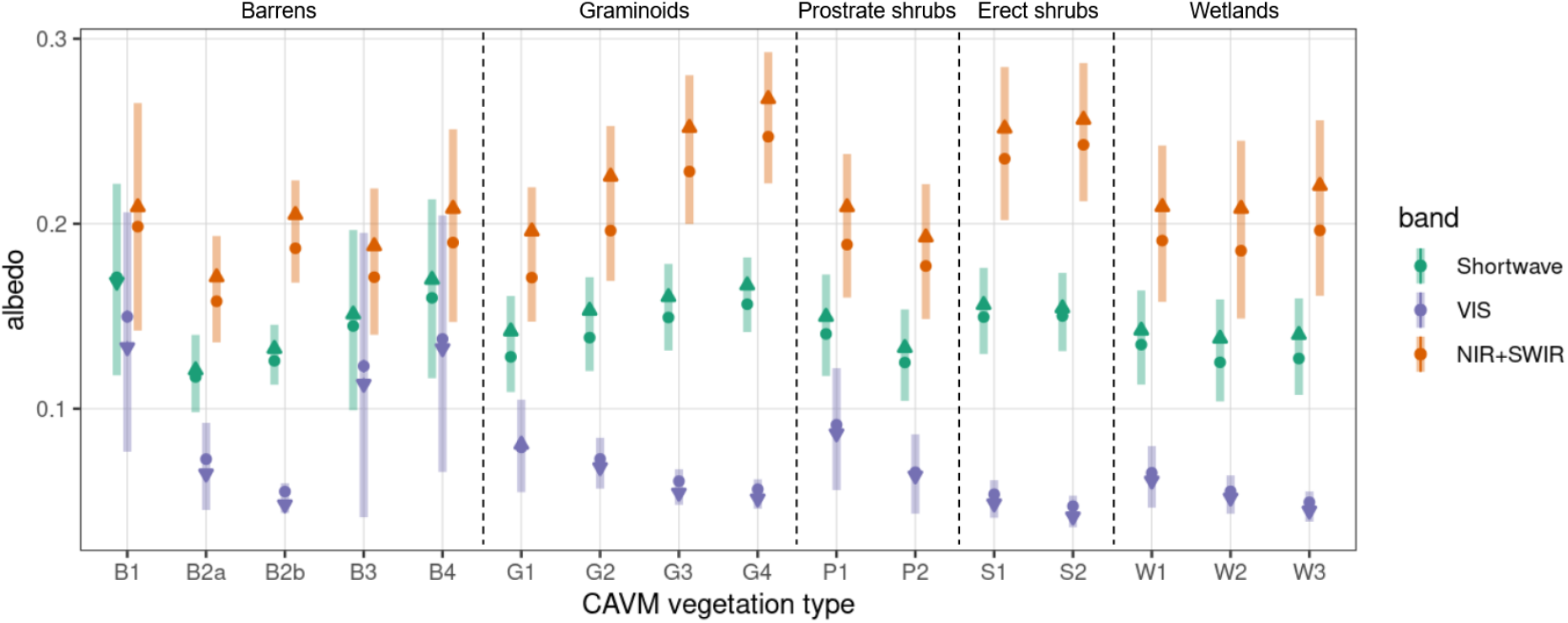
Average albedo of different Arctic vegetation types and the significant changes in albedo over the last 22 years in the broadband shortwave, the VIS, and the NIR+SWIR, spectral domains. The dots represent the average albedo for a given vegetation type at the beginning (2000) and the triangles at the end (2021) of the observed period, based on the trends that show statistical significance (Fig. 1d,e,f). The triangles are pointing up for upward change and down for a downward change. The error bars show the standard deviation of the albedo across the area covered by a vegetation type. The dashed vertical lines separate CAVM vegetation units (B = Barrens, G = Graminoids, P = Prostrate shrubs, S = Erect shrubs, W = Wetlands; whereas 1-4 generally represent an increase in structural complexity (e.g. increasing height) of the canopy; for details see Supplementary Table 1). The data for this figure is available in Supplementary Table 3.

### 2.4. Are those changes reflected in the land surface models?

To answer this question, we compared the shortwave albedo and the albedo change from MODIS observations (Fig. 3, 1st column) to the LSM ensemble mean of 6 CMIP6 climate models (Fig. 3, 2-3rd columns). Due to the restricted time span available for CMIP6 (“land-hist” scenario), the comparison of the two data sets was limited to the period 2000-2014.

**Fig. 3.**
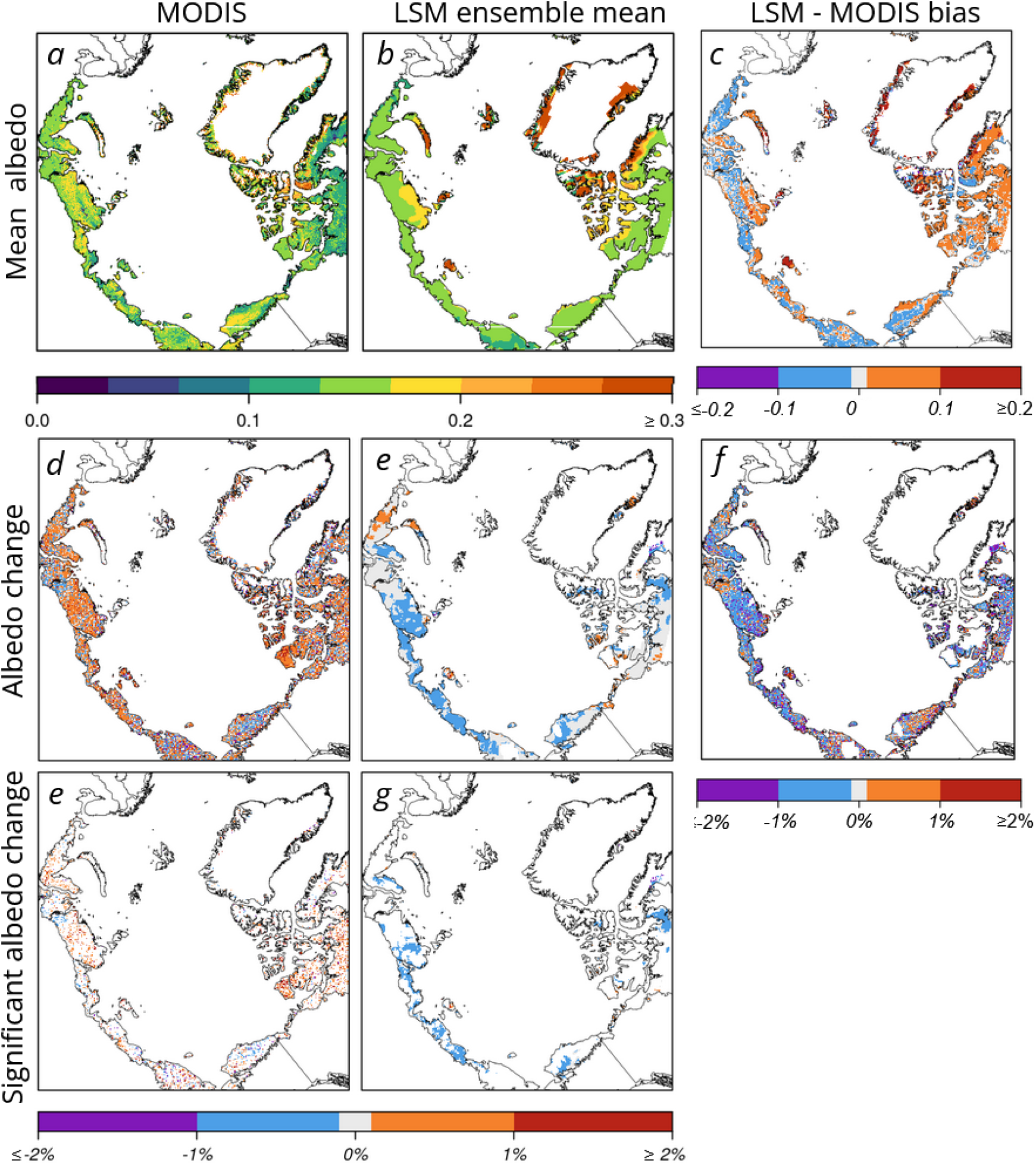
Spatial distribution and comparison of shortwave snow-free mid-summer albedo and its change per year (2000-2014), derived from MODIS and land surface models (LSM), across the Arctic. The plots in the top row depict average values across years of a) MODIS shortwave albedo b) an ensemble mean of 6 LSMs (from CMIP6), and c) the difference between the LSM ensemble mean and MODIS. The next row depicts modeled percent of albedo change per year (based on Sen’s slope) of MODIS (d), LSM ensemble mean (e), and the difference between the two (f). The last row shows significant (p-value < 0.05) albedo change per year for MODIS (e) and LSM ensemble mean (g).

In most of the Russian Arctic area, the land surface model (LSM) ensemble appears to underestimate the MODIS albedo, while over Greenland and Canadian Arctic territories it mostly overestimates the MODIS albedo (Fig. 3a,b,c). The overestimation is especially pronounced in the high Arctic, where the average difference was above 0.1, where the LSM ensemble mean also had large uncertainties (Supplementary Fig. 3a), mostly coming from CNRM-models (Supplementary Fig. 4). The proportion of the area with a decrease in albedo was similar across the LSM ensemble, but much larger than the area derived from MODIS observations (Fig. 3d,e, Supplementary Fig. 3a). The models generally also showed the absence of a significant change over most of the Arctic, but did not locate the significant changes in the same areas as MODIS observations (Fig. 3e,g).

## 3. Discussion

### 3.1. Arctic mid-summer albedo not changing in most areas, and even increasing in some areas

Despite the widely-reported vegetation change and expected corresponding summer albedo decrease (Thompson *et al*., 2004; Chapin *et al*., 2005, Pearson *et al*., 2013), we don’t find significant albedo trends in the shortwave domain in 82% of the pan-Arctic area over the past 22 years (Fig. 1). Areas with significant trends are scattered across the pan-Arctic, and in most cases (14% of area) show an increase in shortwave albedo of 0.1-2% per year since 2000 (Fig. 1). This is in accordance with more recent studies showing a positive average July pan-Arctic trend using AVHRR and MODIS satellite observations (Webb *et al*., 2021, Yu and Leng, 2022).

The stability of shortwave albedo, despite ongoing landscape changes, may be attributed to the fact that the trends are predominantly negative in the VIS domain, but are positive in the NIR+SWIR. Hence, the broadband shortwave trends may be partially canceled out (although only 4% of the area shows a spatial overlap with significant trends in both the VIS and NIR+SWIR albedos).

### 3.2. The mechanisms of increasing albedo

Previous studies have suggested several mechanisms for the unexpected increase in shortwave albedo with assumed biomass increase. They refer to expansion in vegetation and increase in biomass, which affects albedo through a) brighter vegetation replacing darker organic soils (Alessandri *et al*., 2021), b) complex relationships between community-level and leaf-level plant-light interactions (Webb *et al*., 2021) or c) denser vegetation canopies intercepting more snow (Yu and Leng, 2022). However, the sign of the difference between vegetation and soil albedo depends on soil characteristics (e.g. moisture) and other ground cover being overgrown (e.g. mosses and often bright lichens) and there are areas in the lower Arctic not covered by snow in July, showing the same trend direction. So the mechanisms behind the positive biomass-albedo relationship remained puzzling.

Exploring temporal changes in different spectral domains of the shortwave albedo, we find that the VIS domain albedo is indeed decreasing, which is consistent with previous expectations and fulfills the narrative of “darker shrubs replacing lighter vegetation”. However, we also found that 67% of the significant shortwave trends are spatially overlapping with significant NIR+SWIR trends, which implies that shortwave albedo trends are mostly driven by changes in the NIR+SWIR domain. In this domain, vegetation usually reflects and transmits quite large amounts of incoming radiation (Fig. 4). An increase in the biomass and canopy complexity can result in decreased transmittance and increased reflectance (e.g. via increased multiple reflection). Indeed, spectrometer measurements of stacked leaves (Jacquemoud and Ustin, 2019), as well as simulations with increasing leaf area index (Hollinger *et al*., 2010; Wan *et al*., 2021) show this relatively large increase in reflectivity in the NIR+SWIR domain (Fig. 4a,b). In contrast, this effect is much smaller in magnitude in the VIS range, as vegetation absorbs most of the radiation “at the first hit”.

**Fig. 4.**
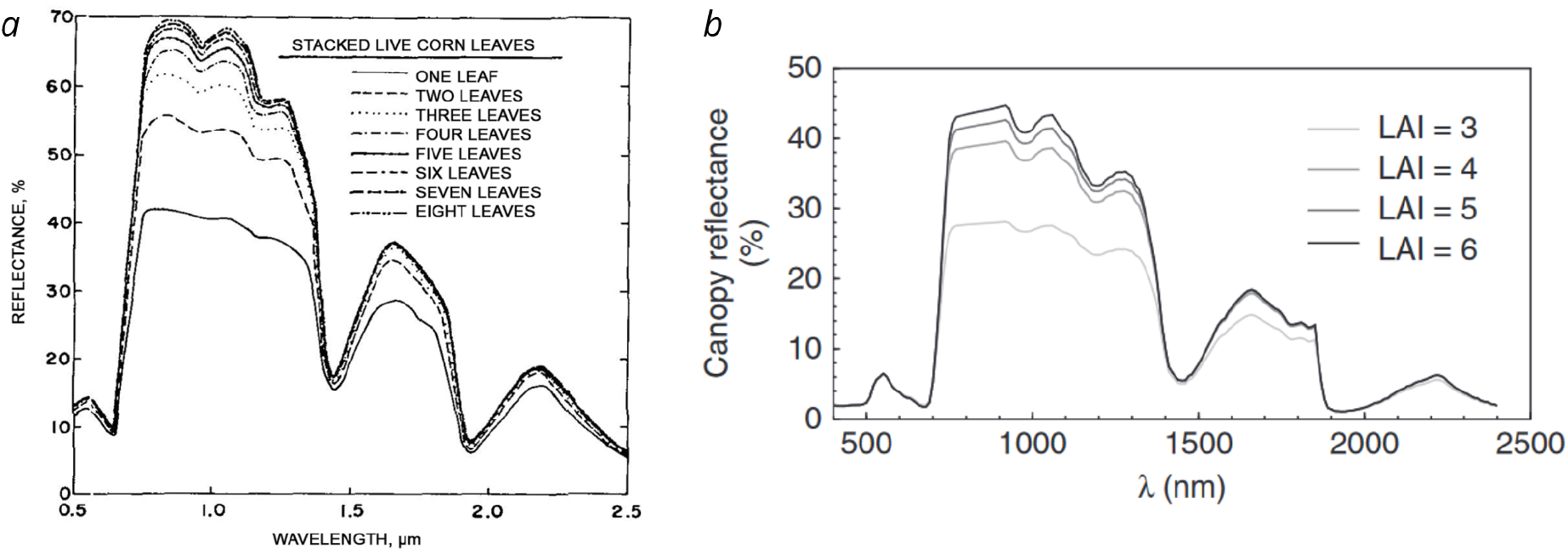
a) Spectral reflectance curves of a) stacked live corn leaves (source: Jacquemoud and Ustin, 2019), b) canopies with different leaf area index (LAI) simulated with the Prospect-SAIL model (source: Hollinger *et al*., 2010).

### 3.3. Albedo change of individual vegetation types and landscape heterogeneity

Despite expected and reported vegetation changes we did not find extensive albedo changes (Fig. 1). This might be explained by spectral effects canceling out (section 3.1) or eventually by the fact that shifts in vegetation types might not be as spatially expansive as expected, e.g. Bjorkman *et al*. (2020) reported most long-term observational studies and warming experiments had no significant shifts in vegetation.

Surprisingly, where albedo change did occur, it was very similar in magnitude, significance and sign across vegetation types (Fig. 2, Supplementary Table 3). For graminoids, we found a curious pattern: along the gradient between G1 (rush/grass, forb, cryptogam tundra) and G4 (tussock-sedge, dwarf-shrub, moss tundra) the difference between VIS and NIR+SWIR albedos becomes larger. We attribute this to the different biomass (e.g. higher shrub abundance and plant height from max. 5 cm in G1 to 40 cm in G4). Erect shrubs also show high differences in average VIS and NIR+SWIR albedo, though the difference between S1 (erect dwarf-shrub tundra) and S2 (low-shrub tundra) types is less pronounced. Barren complexes (≤ 40% plant cover) showed the highest variation in albedo values, which is likely due to residual snow-cover contamination (e.g. underlying frozen ground). We provide detailed data on average albedo and change by each vegetation type (Supplementary Table 3), which can be used for future assessments of how predicted vegetation changes will affect the future Arctic radiative budget, as in Pearson *et al*., (2013).

The Arctic landscape is characterized by very high spatial heterogeneity. Highly patterned permafrost soils influence soil moisture, temperature, and nutrients at meter scales. This creates a fine mosaic of vegetation types, of vegetated and non-vegetated areas (e.g. water cover and bare soil), and of larger topographical features (e.g. river beds, hill slopes, thaw slumps). Vegetation change has been found to be complex and heterogeneous in both satellite-based and plot-based efforts (Bjorkman *et al*., 2020; Myers-Smith *et al*., 2020; CAFF, 2021). For example, Bjorkman *et al*. (2018) found a very widespread increase in height, and increased LAI, but for wet communities only. We also found strong spatial scattering of the areas with significant albedo trends (Fig. 1) which likely relates to the above phenomena. This points to the importance of taking small-scale changes into account. We performed our study on the finest resolution operationally available at pan-Arctic scale and over the last two decades (500m), and recommend that future studies are working on this or higher resolution, and that land surface models advance their resolution from 0.5 degree.

### 3.4. Land surface models do not depict the albedo changes

We found a substantial bias in albedo modeled by the LSM ensemble mean and those observed by the MODIS satellite. This bias is especially pronounced in high latitudes, where the presence of remnant mid-summer snow-cover or permafrost may play a role, especially in the CNRM–CM6 and CNRM-ESM models (Supplementary Fig. 4). This is consistent with previous findings that climate models miss the timing of snow onset and offset, which may result in up to 40% uncertainties in modeling the snow-albedo feedback (Thackeray and Fletcher, 2016). Moreover, the LSMs mostly predicted a decline of shortwave albedo over 2000-2014, whereas MODIS observations show an increase during the same time period (Fig. 3 d,e), and hence, recent trends in albedo are not well captured by the LSMs.

Another explanation for the discrepancy between MODIS observed values and LSM estimates is the limited representation of vegetation types across the terrestrial Arctic in the LSMs. In climate models, vegetation is represented by plant functional types (PFTs), and their distribution and characteristics, together with water and soil fractions, define the modeled albedo. Arctic tundra vegetation is often characterized by only two PFTs (shrub and grass) (Sulman *et al*., 2021), and the plant-light interactions are represented by a simplistic big-leaf model (Bonan *et al*., 2021).

One way to improve climate model estimations is to include additional PFTs such as the CAVM types. Previous studies have shown the importance of detailed vegetation types in land surface energy fluxes in summer and shoulder seasons (Oehri *et al*., in review). Our findings confirm that vegetation type is explaining 22% of the variation in VIS and NIR+SWIR domains and 9% in the broadband shortwave domain (Supplementary Fig. 2). In Supplementary Table 3, we provide, for each vegetation type, the average albedo values and their change per decade across the spectral domains, which can be used for model parameterization. Including the CAVM vegetation types will not only improve the representation of the shortwave energy budget, but also estimates of evapotranspiration, permafrost thawing, and carbon dioxide and methane fluxes.

### 3.5. Study limitations and future directions

Working at the MODIS gridded 500 m spatial resolution limited our study to moderate-scale processes, so we cannot attribute changes in albedo to changes in individual plant traits and their diversity (Juszak *et al*., 2017; Plekhanova *et al*., 2021). This can be further explored by integrating data across scales (Myers-Smith *et al*., 2020). While we mostly focused on vegetation change, other drivers, such as water cover, soil characteristics, and moisture variation all influence albedo. A high-resolution pan-Arctic soil map and temporal soil moisture data, if available, would greatly enhance our understanding of albedo change drivers.

We performed a robust regression analysis, so we were able to detect upward and downward linear trends, but did not test for other shapes of the trend curve. As the p-value threshold to determine significance was chosen as 0.05, we can expect around 5% of the trends to appear significant by chance. Unfortunately, the standard correction of multiple comparisons (Cortés *et al*., 2020) is not applicable here due to a) the heterogeneity of the landscape and the resulting discontinuity of significant trend patterns and b) possible residual snow contamination effects resulting in higher slopes of the trends. Advanced multiple correction methods customized to those limitations would improve the reliability of identified significant trends.

### 3.6. Conclusion

We are the first to provide a pan-Arctic spatially explicit assessment of snow-free mid-summer albedo change separately for broadband shortwave, visible and NIR+SWIR wavelength domains. We show that, in contrast to expectations, the broadband shortwave albedo stayed constant or slightly increased over the past two decades, due to the dominance of the trends in the NIR+SWIR domain, where vegetation and water appear to be key drivers. Thus, widely reported shrubification and increase in NDVI and biomass do not decrease shortwave albedo, as previously expected. We show that land surface models do not incorporate sufficient vegetation-type specific differences and trends in VIS and NIR+SWIR domains which may result in large uncertainties when predicting land-climate feedbacks via albedo. We provide albedo estimates per vegetation type to support a more sophisticated parameterization of climate models, and hence improve predictions of vegetation feedbacks in the Arctic through energy fluxes.

## 4. Methods

### 4.1. MODIS albedo data

Albedo – the fraction of incoming radiation reflected by the surface (unitless).

We used the MODIS (Moderate Resolution Imaging Spectroradiometer) MCD43A3 Albedo product version 6.1 (Schaaf and Wang, 2021). This covers the period from 2000-2021, is retrieved daily, and is based on the best BRDF (bidirectional reflectance distribution function) possible retrieved from 16 days’ worth of inputs with the day of interest emphasized. It has a gridded 500m resolution but can reflect an effective resolution of up to 750m in the mid-latitudes and greater at the poles, affecting the high latitudes (Campagnolo *et al*., 2016). We analyzed white-sky albedo (wholly diffuse bihemispherical reflectance), as it is usually used as input to climate models, which is independent of solar illumination angles. We used VIS, NIR+SWIR, and the broadband shortwave albedo products derived from the first seven spectral land bands of MODIS with narrow-to-broadband conversion coefficients (Wang et. al., 2014). Based on the MCD43A2 quality assessment for each band, we filtered out pixels with QA flag values of 3 (poorer quality magnitude inversion based on fewer than 7 out of 16 observations). In the remaining data, 88% of the pixels had QA flags 0 and 1, indicating best or good quality, full inversion. We used the values for mid-summer (the 15th of July) each year. We further minimized the snow influence by excluding areas with a) snow flag as indicated in the MCD43A2 Snow_BRDF_Albedo band and b) coverage of more than 1% based on the MOD10A2 snow cover product. For our study, we selected the terrestrial Arctic extent based on CAVM treeline (Raynolds *et al*., 2019) and used the Lambert azimuthal equal-area projection.

### 4.2. Trends calculation

To quantify the pan-Arctic albedo trends, we fitted a Mann-Kendall robust line-fit regression (albedo ∼ year) for each MODIS pixel and calculated the slope with Theil–Sen estimator (Sen, 1968) and determined significance with Kendall’s tau statistic (p-value < 0.05). We used the R-package ‘zyp’ for fast Theil-Sen slope calculation (David Bronaugh and Arelia Werner for the Pacific Climate Impacts Consortium, 2019). For the MODIS-derived albedo trend analysis we fitted the regressions over the years 2000-2021, and for the intercomparison with climate models over the years 2000-2014 - as available from the CMIP6 project, land-hist experiment (Eyring *et al*., 2016). We then calculated the percentage of change in albedo per year by dividing the slope by the average albedo value for a given pixel and multiplying by 100.

### 4.3. Vegetation types

We used the new raster version of the Circumpolar Arctic Vegetation Map (CAVM) (Raynolds *et al*., 2019) at a 1 km resolution for determining Arctic vegetation types. In CAVM, vegetation is represented by five broad physiognomic categories: B - barrens, G - graminoid-dominated tundras, P - prostrate-shrub-dominated tundras, S - erect-shrub-dominated tundras, W - wetlands. They are further divided to 16 vegetation types, based on dominant plant functional types (Walker *et al*., 2018; see Supplementary Table 1).

### 4.4. Land surface models

To compare the observed albedo values to the ones currently used in the climate models, we used the 15-year simulation (2000-2014) of 6 CMIP6 project land surface models for which daily values are available: CNRM CM6, CNRM ESM2, IPSL CM6A, MIROC6, UKESM1, EC-Earth3 (Eyring *et al*., 2016). These models operate at 1 degree resolution with global coverage, so we re-gridded them to 0.1 degree resolution and cropped them to the CAVM area. We used 15th of July values and excluded areas covered by snow (“snc” variable larger than 1%). We extracted upward and downward surface radiation at the bottom of the atmosphere (“rsus” and “rsds” variables of land-hist experiment) and calculated the albedo as the fraction of reflected energy. We then extracted the albedo values for 15th of July each year and calculated trends (as in section 4.2) and the average across years. We calculated the model ensemble mean as the average across the 6 models and uncertainty as the standard deviation between the models.

### 4.5. Software

We used Python 3.7.12 (Van Rossum, Guido and Drake, 2009) to download and prepare MODIS data and calculate albedo drivers (Supplementary section 1). The trend analysis, drivers analysis, LSM comparison, and visualizations were done using R 4.1 (R Core Team, 2021) software. The code for main analysis and data for figures are available at the following GitHub repository https://github.com/PlekhanovaElena/aap_paper.

## Acknowledgements

We acknowledge the Service and Support for Science IT team (S3IT, www.s3it.uzh.ch) at the University of Zurich for providing computational servers and technical support. We thank Jeffrey T. Kerby for his comments and questions during study design and article revision. We thank Prof. Dr. Reinhard Furrer for his statistical advice. This study was supported by the University Research Priority Program on Global Change and Biodiversity of the University of Zurich (URPP-GCB) and the Swiss National Science Foundation through project grant no. 178753.

## Authors contributions

EP and GSS designed the study and methods, EP conducted the analyses, JSK pre-processed climate data, EP, GSS, and JSK drafted the article (text & figures), all authors (EP, JO, JSK, AE, CS, GSS) discussed the results and revised the article.

