## Supplementary Materials for "Mid-summer snow-free albedo across the Arctic tundra was mostly stable or increased over the past two decades"

**Supplementary Table 1**. Description of the Circumpolar Arctic Vegetation map (CAVM) vegetation types as in Walker *et al.*, (2010).

| CAVM category | type | Description |
|---|---|---|
| Barrens | B1 - Cryptogam, herb barren | Dry to wet barren landscapes with very sparse, very low-growing plant cover. Scattered herbs, lichens, mosses, and liverworts. |
| | B2a,b - Cryptogam barren complex (bedrock) | Areas of exposed rock and lichens interspersed with lakes and more vegetated areas, as found on the Canadian Shield. |
| | B3 - Noncarbonate mountain complex | Mountain vegetation on noncarbonate bedrock. The variety and size of plants decrease with elevation and latitude. Hatching colour and code indicate the bioclimate subzone at the mountain base. |
| | B4 - Carbonate mountain complex | Mountain vegetation on carbonate bedrock. The variety and size of plants decrease with elevation and latitude. Hatching colour and code indicate the bioclimate subzone at the mountain base. |
| Graminoids | G1 - Rush/grass, forb, cryptogam tundra | Moist tundra with moderate to complete cover of very low-growing plants. Mostly grasses, rushes, forbs, mosses, lichens, and liverworts. |
| | G2 - Graminoid, prostrate dwarf-shrub, forb tundra | Moist to dry tundra, with open to continuous plant cover. Sedges are dominant, along with prostrate shrubs <5 cm tall. |
| | G3 - Non-tussock sedge, dwarf-shrub, moss tundra | Moist tundra dominated by sedges and dwarf shrubs <40 cm tall, with well-developed moss layer. Barren patches due to frost boils and periglacial features are common. |
| | G4 - Tussock-sedge, dwarf-shrub, moss tundra | Moist tundra, dominated by tussock cottongrass (*Eriophorum vaginatum*) and dwarf shrubs <40 cm tall. Mosses are abundant. |
| Prostrate shrubs | P1 - Prostrate dwarf-shrub, herb tundra | Dry tundra with patchy vegetation. Prostrate shrubs <5 cm tall (such as Dryas and Salix arctica) are dominant, with graminoids and forbs. Lichens are also common. |
| | P2 - Prostrate/hemiprostrate dwarf-shrub tundra | Moist to dry tundra dominated by prostrate and hemiprostrate shrubs <15 cm tall, particularly Cassiope. |
| Erect shrubs | S1 - Erect dwarf-shrub tundra | Tundra dominated by erect dwarf-shrubs, mostly <40 cm tall. |
| | S2 - Low-shrub tundra | Tundra dominated by low shrubs >40 cm tall. |
| Wetlands | W1 - Sedge/grass, | Wetland complexes in the colder areas of the |

| | | |
|---|---|---|
| | moss wetland | Arctic, dominated by sedges, grasses, and mosses. |
| | W2 - Sedge, moss, dwarf-shrub wetland | Wetland complexes in the milder areas of the Arctic, dominated by sedges, grasses, and mosses, but including dwarf shrubs <40 cm tall. |
| | W3 - Sedge, moss, low-shrub wetland | Wetland complexes in the warmer areas of the Arctic, dominated by sedges and low shrubs >40 cm tall. |

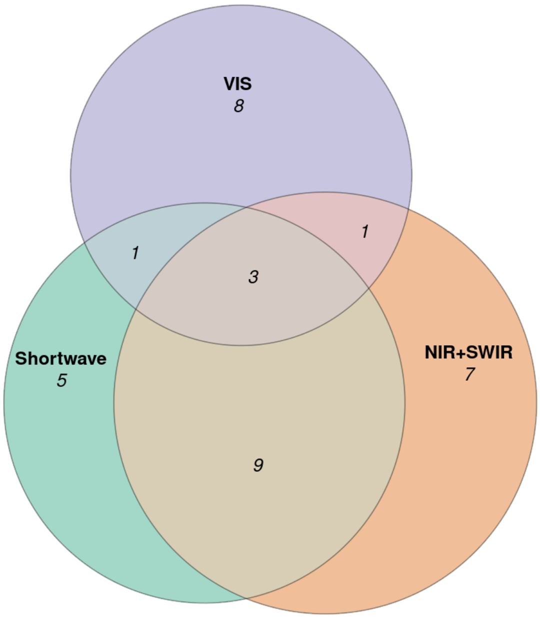


**Supplementary Fig. 1.** Euler diagram showing overlap between significant albedo trends in shortwave (SW), VIS and NIR domains (see Fig. 1). Each number is a percentage from total trends - e.g. in VIS range (violet circle), 13% of trends were significant, from them 4% overlapped with significant NIR trends, 4% overlapped with significant shortwave and 3% overlapped with both.

---

### ***Supplementary section 1.*** *Drivers analysis of snow-free albedo across space*

What drives the observed change? Using space for time substitution, we investigated how land surface properties, phenological and climatic factors influence snow-free mid-summer albedo averaged across years 2000-2021 (see Supplementary Fig.2).

To estimate how different factors influence albedo across space, we fitted a linear regression over land cover, climatic and phenological factors on the 100,000 randomly chosen pixels. We calculated water cover based on superfine water index (Sharma *et al.*, 2015), derived climatic factors and first growing day (fgs) from the CHELSA database (Karger *et al.*, 2017) and averaged those predictors over the years (see Supplementary Table 2 for more details). Using the lmg function of the R-package „relaimpo“ (Grömping, 2007), we calculated variance partitioning based on $R^2$ contribution averaged over orderings among the regressors.

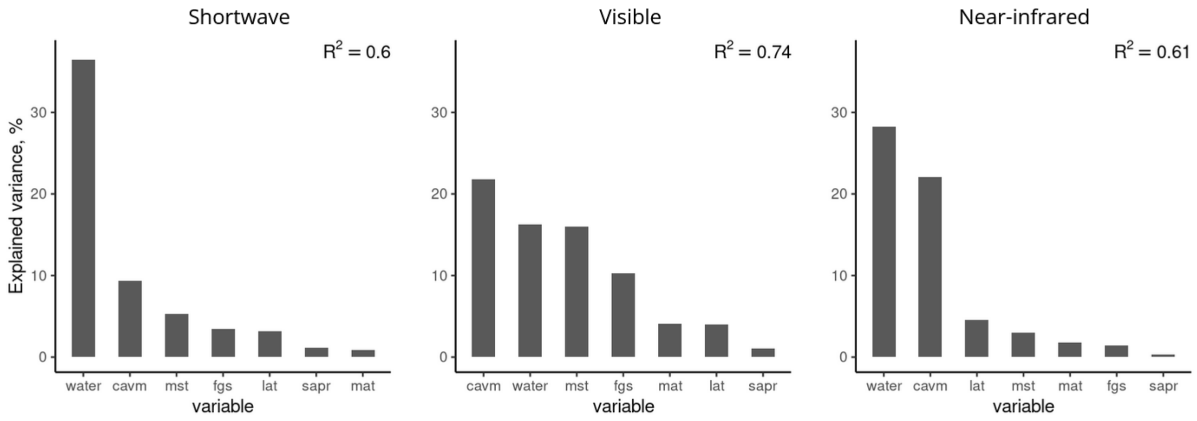


**Supplementary Fig. 2.** Percentage of shortwave, VIS and NIR albedo spatial variance explained by different factors. The explained variance was calculated as the $R^2$ contribution averaged over orderings among regressors. The factors are the following: Water - water fraction, CAVM - vegetation type, MST - mean summer temperature, FGS - first day of the growing season, lat - latitude, SAPR - sum of annual precipitation, MAT - mean annual temperature, LAT - latitude.

The chosen predictors explained 60% of shortwave albedo variance, 74% of VIS and 61% of NIR albedo. Water fraction was the leading factor in shortwave and NIR domains, explaining 36% and 28% of albedo correspondingly. CAVM type was a dominant factor for VIS domain (22%), second-most important for NIR (22%) and shortwave (9%) domains. Mean summer temperature and first growing season day were mostly influencing the VIS domain (17% and 10%).

**Supplementary Table 2**. Predictors used in the albedo drivers analysis. All data sources are standard, validated, open source products.

| Variable | Source | Details |
|---|---|---|
| Albedo | MCD43A3 V006/V061 (Schaaf and Wang, 2015) | Albedo_WSA_Shortwave, Albedo_WSA_vis, Albedo_WSA_nir |
| CAVM type | Raster version of CAVM (Raynolds *et al.*, 2019) | All the CAVM types, except water and glaciers |
| Mean annual temperature | CHELSA v.1.2 (Karger *et al.*, 2017) | Calculated from monthly *tas* variable for 2000-2019 |
| Mean summer | CHELSA v.1.2 (Karger *et al.*, 2017) | Calculated from monthly *tas* |

| | | |
|---|---|---|
| temperature | | variable for 2000-2019 |
| Sum of annual precipitation | CHELSA v.1.2 (Karger *et al.*, 2017) | Calculated from monthly *pr* variable for 2000-2019 |
| First day of the growing season | CHELSA v.2.1 (Karger *et al.*, 2017) | *fgd* variable |
| Water cover | MCD43A4 V006/V061 (Schaaf and Wang, 2021) | Water cover for each pixel was calculated based on superfine index (Sharma *et al.*, 2015) |

---

**Supplementary Table 3**. Statistics of albedo values, significant slopes and changes over the 2000-2022 period for each CAVM vegetation type (Walker *et al.*, 2010). This table also available as a separate CSV document attached to the electronic version of this manuscript.

| band | CAVM type | Mean albedo value | Std albedo value | Mean significant slope | Std significant slope | % of significant slopes | % of positive slopes | Mean change over 22 yrs |
|---|---|---|---|---|---|---|---|---|
| shortwave | B1 | 0.169834 | 0.05178 | -0.00012 | 0.002586 | 13 | 57 | -0.0027 |
| shortwave | B2a | 0.119021 | 0.020879 | 0.000185 | 0.001012 | 18 | 71 | 0.0041 |
| shortwave | B2b | 0.129171 | 0.016183 | 0.000329 | 0.000813 | 21 | 80 | 0.0072 |
| shortwave | B3 | 0.147899 | 0.048725 | 0.000316 | 0.002554 | 14 | 76 | 0.007 |
| shortwave | B4 | 0.164893 | 0.04839 | 0.000495 | 0.001872 | 11 | 73 | 0.0109 |
| shortwave | G1 | 0.134986 | 0.025978 | 0.000695 | 0.00122 | 21 | 84 | 0.0153 |
| shortwave | G2 | 0.145756 | 0.025382 | 0.000731 | 0.000989 | 24 | 85 | 0.0161 |
| shortwave | G3 | 0.154898 | 0.023452 | 0.000556 | 0.000802 | 25 | 84 | 0.0122 |
| shortwave | G4 | 0.161651 | 0.020166 | 0.000514 | 0.00092 | 20 | 82 | 0.0113 |
| shortwave | P1 | 0.145122 | 0.027447 | 0.000467 | 0.001253 | 20 | 76 | 0.0103 |
| shortwave | P2 | 0.128964 | 0.024694 | 0.000398 | 0.001037 | 21 | 74 | 0.0088 |
| shortwave | S1 | 0.15288 | 0.023266 | 0.00033 | 0.000872 | 19 | 72 | 0.0073 |
| shortwave | S2 | 0.1523 | 0.021166 | 0.000212 | 0.000868 | 15 | 66 | 0.0047 |
| shortwave | W1 | 0.138506 | 0.025485 | 0.000385 | 0.001484 | 22 | 69 | 0.0085 |
| shortwave | W2 | 0.131545 | 0.027584 | 0.000646 | 0.001229 | 29 | 80 | 0.0142 |
| shortwave | W3 | 0.1336 | 0.026078 | 0.000642 | 0.001029 | 20 | 84 | 0.0141 |
| vis | B1 | 0.141481 | 0.064697 | -0.00083 | 0.003444 | 11 | 37 | -0.0182 |

| | | | | | | | | |
|---|---|---|---|---|---|---|---|---|
| vis | B2a | 0.068821 | 0.023596 | -0.00039 | 0.001169 | 15 | 18 | -0.0087 |
| vis | B2b | 0.051715 | 0.008 | -0.00035 | 0.000262 | 18 | 4 | -0.0077 |
| vis | B3 | 0.118216 | 0.076854 | -4.80E-04 | 0.00343 | 11 | 39 | -1.07E-02 |
| vis | B4 | 0.135176 | 0.069373 | -2.50E-04 | 0.002581 | 10 | 39 | -0.0056 |
| vis | G1 | 0.079896 | 0.02496 | 8.00E-05 | 0.00149 | 15 | 61 | 0.0018 |
| vis | G2 | 0.07059 | 0.013778 | -0.00023 | 0.000903 | 15 | 34 | -0.005 |
| vis | G3 | 0.057686 | 0.009648 | -0.00032 | 0.00041 | 14 | 15 | -0.007 |
| vis | G4 | 0.053936 | 0.007938 | -0.00025 | 0.000419 | 16 | 19 | -0.0056 |
| vis | P1 | 0.08897 | 0.032953 | -0.00024 | 0.001518 | 15 | 38 | -0.0052 |
| vis | P2 | 0.064724 | 0.021436 | -9.01E-05 | 0.001041 | 15 | 41 | -0.002 |
| vis | S1 | 0.051242 | 0.010168 | -0.00025 | 0.000418 | 15 | 17 | -0.0055 |
| vis | S2 | 0.044505 | 0.008537 | -0.00028 | 0.000317 | 14 | 9 | -0.0062 |
| vis | W1 | 0.063193 | 0.016601 | -0.00021 | 0.00088 | 18 | 36 | -0.0047 |
| vis | W2 | 0.053613 | 0.010352 | -0.00018 | 0.000653 | 16 | 36 | -0.0039 |
| vis | W3 | 0.047131 | 0.008193 | -0.00023 | 0.000478 | 14 | 24 | -0.0051 |
| nir+swir | B1 | 0.203747 | 0.061464 | 0.000529 | 0.002745 | 12 | 68 | 0.0116 |
| nir+swir | B2a | 0.164632 | 0.028752 | 0.000653 | 0.001179 | 21 | 85 | 0.0144 |
| nir+swir | B2b | 0.195799 | 0.027671 | 0.000906 | 0.000847 | 24 | 95 | 0.0199 |
| nir+swir | B3 | 0.179542 | 0.039568 | 0.000842 | 0.001976 | 14 | 79 | 0.0185 |
| nir+swir | B4 | 0.199017 | 0.052073 | 0.000918 | 0.001974 | 12 | 80 | 0.0202 |
| nir+swir | G1 | 0.183426 | 0.036281 | 0.001253 | 0.001733 | 19 | 86 | 0.0276 |
| nir+swir | G2 | 0.210946 | 0.041889 | 0.001466 | 0.001456 | 23 | 91 | 0.0322 |
| nir+swir | G3 | 0.240094 | 0.040205 | 0.001185 | 0.001299 | 25 | 90 | 0.0261 |
| nir+swir | G4 | 0.257298 | 0.035491 | 0.001024 | 0.001553 | 22 | 86 | 0.0225 |
| nir+swir | P1 | 0.198889 | 0.038803 | 0.001018 | 0.001537 | 19 | 84 | 0.0224 |
| nir+swir | P2 | 0.184963 | 0.036491 | 0.000779 | 0.001574 | 22 | 77 | 0.0171 |
| nir+swir | S1 | 0.243354 | 0.041379 | 0.000822 | 0.001409 | 18 | 80 | 0.0181 |
| nir+swir | S2 | 0.249512 | 0.037386 | 0.000682 | 0.001428 | 14 | 77 | 0.015 |
| nir+swir | W1 | 0.200018 | 0.042271 | 0.000912 | 0.002464 | 22 | 73 | 0.0201 |
| nir+swir | W2 | 0.196817 | 0.048075 | 0.001139 | 0.002113 | 29 | 80 | 0.0251 |
| nir+swir | W3 | 0.208502 | 0.047465 | 0.00121 | 0.001752 | 20 | 86 | 0.0266 |

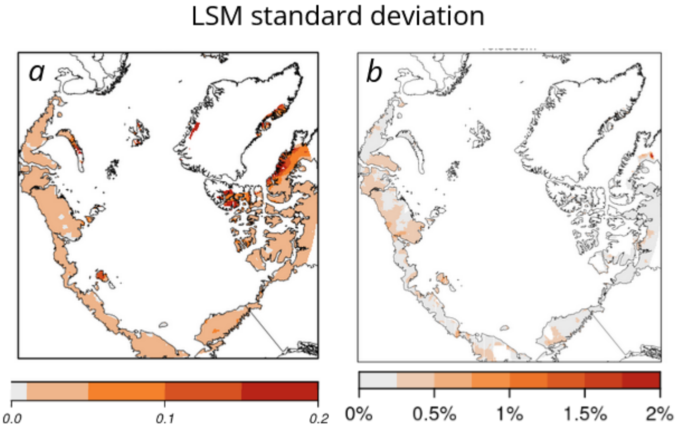


**Supplementary Figure 3.** Spatial distribution of standard deviation of a) Mid-summer snow-free shortwave albedo modelled by LSM ensemble and averaged across 2000-2014 and b) its change per year (2000-2014), over the Arctic. The values below 0.01 (a) and albedo change below 0.1% per year (b) is shown in grey.

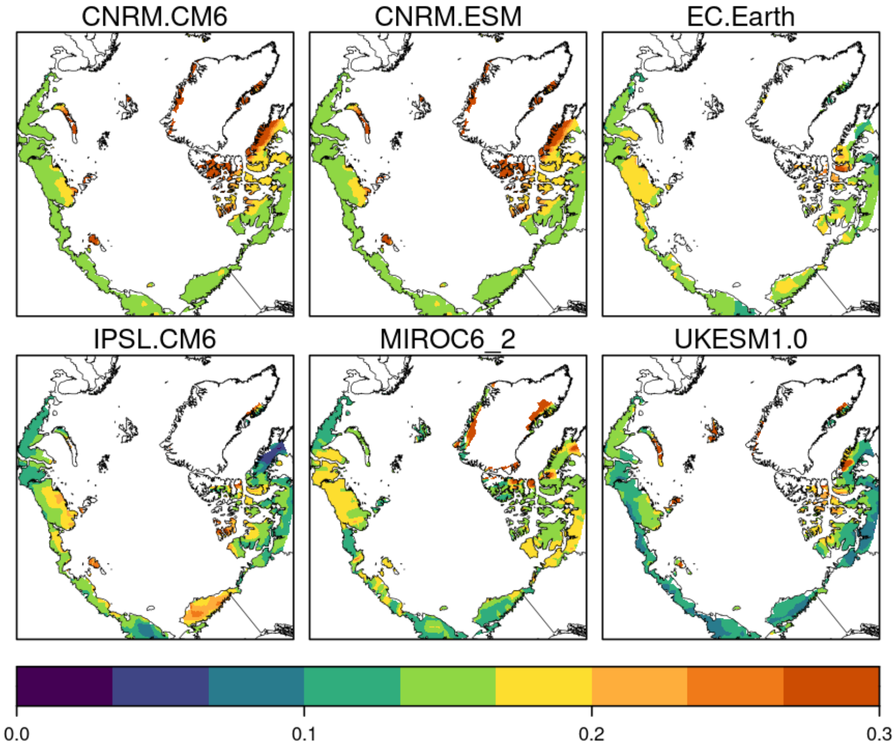


**Supplementary Figure 4.** Spatial distribution of average (across years 2000-2014) July shortwave albedo values of 6 LSM (from CMIP6) used in this study.

***Supplementary section 2.*** *MODIS data quality assessment*

We used MCD43A2 QA product (BRDF_Albedo_Band_Quality_BandN) to assess the quality of individual bands.

**Quality info**

0 = best quality, full inversion (WoDs, RMSE majority good)
1 = good quality, full inversion (also including the cases that no clear sky observations over the day of interest or the Solar Zenith Angle is too large even WoDs, RMSE majority good)
2 = Magnitude inversion (numobs >=7)
3 = Magnitude inversion (numobs >=2&<7)
4 = Fill value

**Bands info**

| Band | spectral range [nm] | Description |
|---|---|---|
| 1 | 620–670 | red |
| 2 | 841–876 | NIR |
| 3 | 459–479 | blue |
| 4 | 545–565 | green |
| 5 | 1230–1250 | SWIR 1 |
| 6 | 1628–1652 | SWIR 2 |
| 7 | 2105–2155 | SWIR 3 |

**Quality filtering**

We treat
QA = 0,1,2 as QA = 0 - good quality, and
QA =3,4 as QA = 1 - bad quality.

For the VIS range we show on the picture on the right bands 1,3,4. 6-22% of values have bad quality (QA = 1 flag). For the NIR range we analysed bands 2,5,6,7. Similarly, 6-21% of values had QA = 1 flag. We excluded all the QA=1 pixels from the analysis. The excluded pixels were differently spatially distributed across years (Supplementary Figure 5).

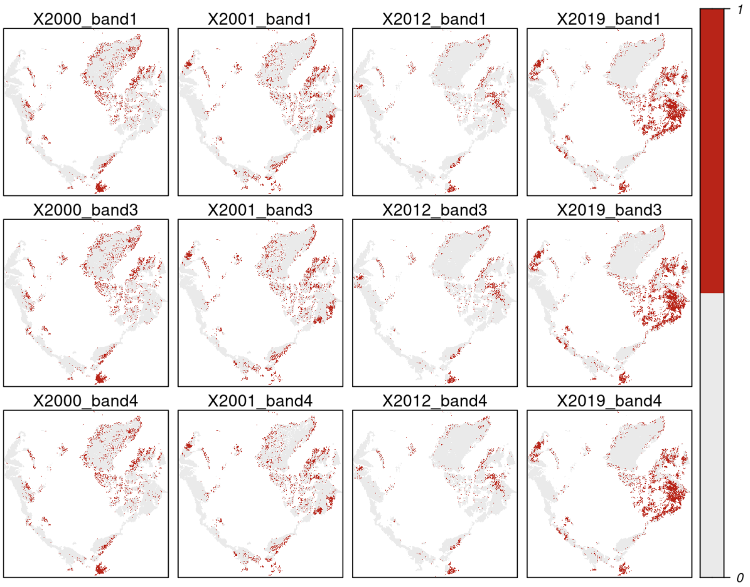


**Supplementary Figure 5.** An example of spatial distribution of quality flags for bands 1,3 and 4 for years 2000, 2001, 2012 and 2019.
